# SAINT: Self-Attention Augmented Inception-Inside-Inception Network Improves Protein Secondary Structure Prediction

**DOI:** 10.1101/786921

**Authors:** Mostofa Rafid Uddin, Sazan Mahbub, M Saifur Rahman, Md Shamsuzzoha Bayzid

## Abstract

**Motivation:** Protein structures provide basic insight into how they can interact with other proteins, their functions and biological roles in an organism. Experimental methods (e.g., X-ray crystallography, nuclear magnetic resonance spectroscopy) for predicting the secondary structure (SS) of proteins are very expensive and time consuming. Therefore, developing efficient computational approaches for predicting the secondary structure of protein is of utmost importance. Advances in developing highly accurate SS prediction methods have mostly been focused on 3-class (Q3) structure prediction. However, 8-class (Q8) resolution of secondary structure contains more useful information and is much more challenging than the Q3 prediction.

**Results:** We present SAINT, a highly accurate method for Q8 structure prediction, which incorporates self-attention mechanism (a concept from natural language processing) with the Deep Inception-Inside-Inception (Deep3I) network in order to effectively capture both the *short-range* and *long-range interactions* among the amino acid residues. SAINT offers a more interpretable framework than the typical black-box deep neural network methods. Through an extensive evaluation study, we report the performance of SAINT in comparison with the existing best methods on a collection of benchmark datasets, namely, TEST2016, TEST2018, CASP12 and CASP13. Our results suggest that self-attention mechanism improves the prediction accuracy and outperforms the existing best alternate methods. SAINT is the first of its kind and offers the best known Q8 accuracy. Thus, we believe SAINT represents a major step towards the accurate and reliable prediction of secondary structures of proteins.

**Availability:** SAINT is freely available as an open source project at https://github.com/SAINTProtein/SAINT.

## 1 Introduction

Proteins are bio-molecules made of long chains of amino acid residues connected by peptide bonds. The functions of proteins are usually determined by the their tertiary structure and for determining the tertiary structure and related properties, the secondary structure information is crucial. Protein structure can be experimentally determined by X-ray crystallography and multi-dimensional magnetic resonance in laboratory, but these methods are very costly and time consuming and are yet to be consistent with the proliferation of protein sequence data [1]. Thus, the proteins with known primary sequence continue to outnumber the proteins with experimentally determined secondary structures. The structural properties of a protein depend on its primary sequence [2–5], yet it remains as a difficult task to accurately determine the secondary and tertiary structures of proteins. Hence, the problem of predicting the structures of a protein – given its primary sequence – is crucially important and remains as one of the greatest challenges in computational biology.

Secondary structure – a conformation of the local structure of the polypeptide backbone – prediction dates back to the work of Pauling and Corey in 1951 [6]. The secondary structures of proteins are traditionally characterized as 3 states (Q3): helix (H), strand (E), and coil (C). Afterwards, a more fine-grained characterization of the secondary structures was proposed [7] for more precise information by extending the three states into eight states (Q8): *α*-helix (H), 3_10_-helix (G), *π*-helix (I), *β*-strand (E), isolated *β*-bridge (B), turn (T), bend (S), and Others (C). Q8 prediction is more challenging and can reveal more precise and high resolution on the structural properties of proteins.

Protein secondary structure prediction is an extensively studied field of research [8–30]. Developing computational approaches (especially using machine learning techniques) for 3-state SS prediction has a long history which dates back to the works of Qian & Sejnowski [8] and Holley & Karplus [9] who first used neural networks to predict SS. In the 1980s, only statistical model based methods were used on raw sequence data which could ensure Q3 accuracy merely below 60%. Afterwards, significant improvement was achieved [10–12] by leveraging the evolutionary information such as the position-specific score matrices (PSSM) derived from multiple sequence alignments. Subsequently, many machine learning methods have been developed for Q3 prediction which include support vector machines (SVM) [13–15,31], probabilistic graphical models [16,32,33], hidden Markov models [17,18], bidirectional recurrent neural networks [19–22,34,35], and other deep learning frameworks [23, 36].

The performance of Q3 prediction methods has approached the postulated theoretical limit [24]. At the same time, there has now been a growing awareness that 8-state prediction can reveal more valuable structural properties. Accurate 8-state secondary structures predictions can reduce the search space in template-free protein tertiary structure modeling by restricting the variations of backbone dihedral angles within a small range according to the Ramachandran plots [37,38]. Also, differentiation among 3_10_ helix, *α*-helix, and *π*-helix in secondary structure prediction helps to assign residues and fit protein structure models in cryo-electron microscopy density maps [38, 39]. As such, the interest of the research community has recently shifted from Q3 prediction to relatively more challenging Q8 prediction. Quite a few deep learning methods for Q8 prediction have been proposed over the last few years [19,25,26,28–30,40]. To the best of our knowledge, the first notable success in Q8 prediction methods was SSpro8 [19] which was published in 2002 and achieved 63.5% Q8 accuracy on the benchmark CB513 dataset [41], 64.9% on CASP10 and 65.6% on CASP11 [25]. Later in 2011, RaptorX-SS8 [40], another 8 state predictor using conditional neural fields, surpassed SSpro8 by demonstrating 64.9% Q8 accuracy on CB513. In 2014, Zhou and Troyanskaya [26] highlighted the challenges in 8-state prediction and obtained 66.4% Q8 accuracy on CB513 dataset using deep generative stochastic network (GSN). Some of the notable subsequent works include deep conditional random fields (DeepCNF) [25], cascaded convolutional and recurrent neural network (DCRNN) [27], next-step conditioned deep convolutional neural network (NC-CNN) [28], multi-scale CNN with highway (CNNH_PSS) [29], DeepACLSTM [42] with an asymmetric convolutional neural networks (ACNNs) combined with bidirectional long short-term memory (BLSTM), deep inception-inside-inception (Deep3I) network named MUFOLD-SS [30], CNN and Bidirectional LSTM based network NetSurfP-2.0 [43], and SPOT-1D [44] which is an ensemble of of hybrid models consisting of Residual Convolutional Neural Networks (ResNet) and 2-Dimensional Bidirectional Residual LSTM Networks (2D-BRLSTM). While most of the methods use sequence data and sequence profiles obtained from Position Specific Scoring Matrix (PSSM) as features, the more recent methods, such as, MUFOLD-SS, CRRNN, NetSurfP-2.0, and SPOT-1D leveraged HMM profiles and physicochemical properties of residues as well. The most recent and accurate method SPOT-1D [44] also used predicted contact map information as features and could achieve a significant boost in accuracy. Although these works demonstrate a steady improvement in the published Q8 accuracy over the past few years, the improvements across successive publications are very small. Nevertheless, these small improvements are considered significant given the high complexity of 8-state SS prediction.

Usually the models that focus more on short range dependencies (local context of the amino acid residues) face difficulties in effectively capturing the long range dependencies (interactions between amino acid residues that are close in three-dimensional space, but far from each other in the primary sequence) [22,27,45]. Various deep learning based models have been leveraged to handle the long-range interactions by using recurrent or highway networks [28,29], deeper networks with convolutional blocks [30], long short-term memory (LSTM) cells [22,27], whereas the short-range interactions have been handled by convolutional blocks of smaller window size [27, 28, 30]. These methods circumvent some challenging issues in capturing the non-local interactions, but have limitations of their own. Models, using recurrent neural networks to capture long range dependencies, may suffer from *vanishing gradient* or *exploding gradient* problems [46–49]. Moreover, these methods may fail to effectively capture the dependencies when the sequences are very long [50]. Furthermore, as the models grow deeper, the number of parameters also grows which makes it prone to over-fitting. It is also likely that the short range relationships captured in the earlier (shallow) layers may disappear as the models grow deeper [29]. As a result, developing techniques which can capture both long-range and short-range dependencies simultaneously is of utmost importance. Another limiting factor of the deep learning methods is that the high accuracy comes at the expense of high abstraction (less interpretability) due to their black-box nature [51–54]. Although there has been a flurry of recent works towards designing deep learning techniques for bio-molecular data, no notable attempt has been made in developing methods with improved interpretablity and explainability – models that are able to summarize the reasons of the network behavior, or produce insights about the causes of their decisions and thus gain trust of users.

In this study, we present SAINT (**S**elf-**A**ttention Augmented **I**nception Inside Inception **N**e**T**work) – a novel method for 8-state SS prediction which uniquely incorporates the *self-attention mechanism* [55] with a state-of-the-art Deep Inception-Inside-Inception (Deep3I) network [30]. We proposed a novel architecture called attention-augmented 3I (2A3I) in order to capture both the local- and long-range interactions. SAINT was compared with a collection of the best alternate methods for Q8 prediction on CASP12 and CASP13 as well as on more recent, challenging and larger test sets (*TEST2016* and *TEST2018*), that were analyzed by a recent and highly accurate method SPOT-1D [44]. SAINT obtained superior Q8 accuracy compared to state-of-the-art predictors on the benchmark datasets – 77.73% accuracy on TEST2016, 76.09% on TEST2018, 74.78% on CASP13, 74.17% in CASP12, and 72.25% on the CASP Free Modeling (FM) targets. SAINT also obtained high precision, recall and *F*1-score for individual states. Moreover, SAINT provides interesting insights regarding the interactions and roles of amino acid residues while forming secondary structures, which help to interpret how the predictions are made. Thus, we have made the following significant contributions: 1) we, for the first time, successfully translated the success of self-attention mechanism from natural language processing to the domain of protein structure prediction, and demonstrated that self-attention improves the accuracy SS prediction, 2) introduced a method which can capture both the short- and long-range dependencies, and offers the best known Q8 accuracy, and 3) improved the interpretability of the black-box deep neural network based methods which are often criticized for lack of interpretability.

## 2 Approach

### 2.1 Feature Representation

SAINT takes a protein sequence feature vector *X* = (*x*_1_, *x*_2_, *x*_3_,…, *x_N_*) as input, where *x_i_* is the vector corresponding to the *i^th^* residue, and it returns the protein structure label sequence vector *Y* = (*y*_1_, *y*_2_, *y*_3_,…, *y_N_*) as output, where *y_i_* is the structure label (one of the eight possible states) of the *i^th^* residue. Similar to SPOT-1D-base and MUFOLD-SS, our base model contains 57 features from PSSM profiles, HHM profiles and physicochemical properties. To generate PSSM, PSI-BLAST [56] was run against Uniref90 database [57] with inclusion threshold 0.001 and three iterations. The HHM profiles were generated using HHblits [58] using default parameters against uniprot20_2013_03 sequence database, which can be downloaded from http://www.user.gwdg.de/~compbiol/data/hhsuite/databases/hhsuite_dbs/. HHblits also generates 7 transition probabilities and 3 local alignment diversity values which we used as features as well. Seven physicochemical properties of each amino acid (e.g., steric parameters (graph-shape index), polarizability, normalized van der Waals volume, hydrophobicity, isoelectric point, helix probability, and sheet probability) were obtained from Meiler *et al*. [59]. So, in our base model, the dimension of *x_i_* is 57 as this is the concatenation of 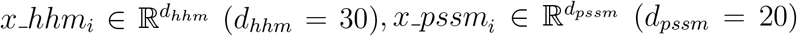, and 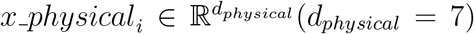. Additional features were generated by windowing the predicted contact information as was done in SPOT-1D. The contact maps were generated using SPOT-contact [60] locally and were used as our features by varying window lengths(the number of preceding or succeeding residues whose pairwise contact information were extracted for a target residue). Our ensemble model constitutes of four different models, that we trained with varying input features: one without the contact maps (base model) and three with different window lengths (10,20, and 50) of the contact-map-based features. The features were normalized to ensure 0 mean and standard deviation of 1 in the training data, similar to SPOT-1D.

### 2.2 Architecture of SAINT

The architecture of SAINT can be split into three separate discussions: 1) the architecture of our proposed self-attention module, 2) the architecture of the existing inception module and the proposed attention augmented inception module, and finally 3) the overall pipeline of SAINT.

#### 2.2.1 Self-attention module

Attention mechanism implies paying attention to specific parts of input data or features while generating output sequence [55,61]. It calculates a probability distribution over the elements in the input sequence and then takes the weighted sum of those elements based on this probability distribution while generating outputs.

In self-attention mechanism [55, 62, 63], each vector in the input sequence is transformed into three vectors-*query, key* and *value*, by three different functions. Each of the output vectors is a weighted sum of the *value* vectors, where the weights are calculated based on the compatibility of the *query* vectors with the *key* vectors by a special function, called *compatibility function* (discussed later in this section).

The self-attention module we designed and augmented with the Deep3I network [30] is inspired from the self-attention module proposed by Vaswani *et al*. [55] and is depicted in Fig. 1. Our self-attention module takes two inputs: 1) the features from the previous inception module or layer, 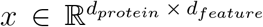, and 2) position identifiers, *pos-id* ∈ 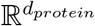, where *d_protein_* is the length of the protein sequence, and *d_feature_* is the length of the feature vector.

**Figure 1:**
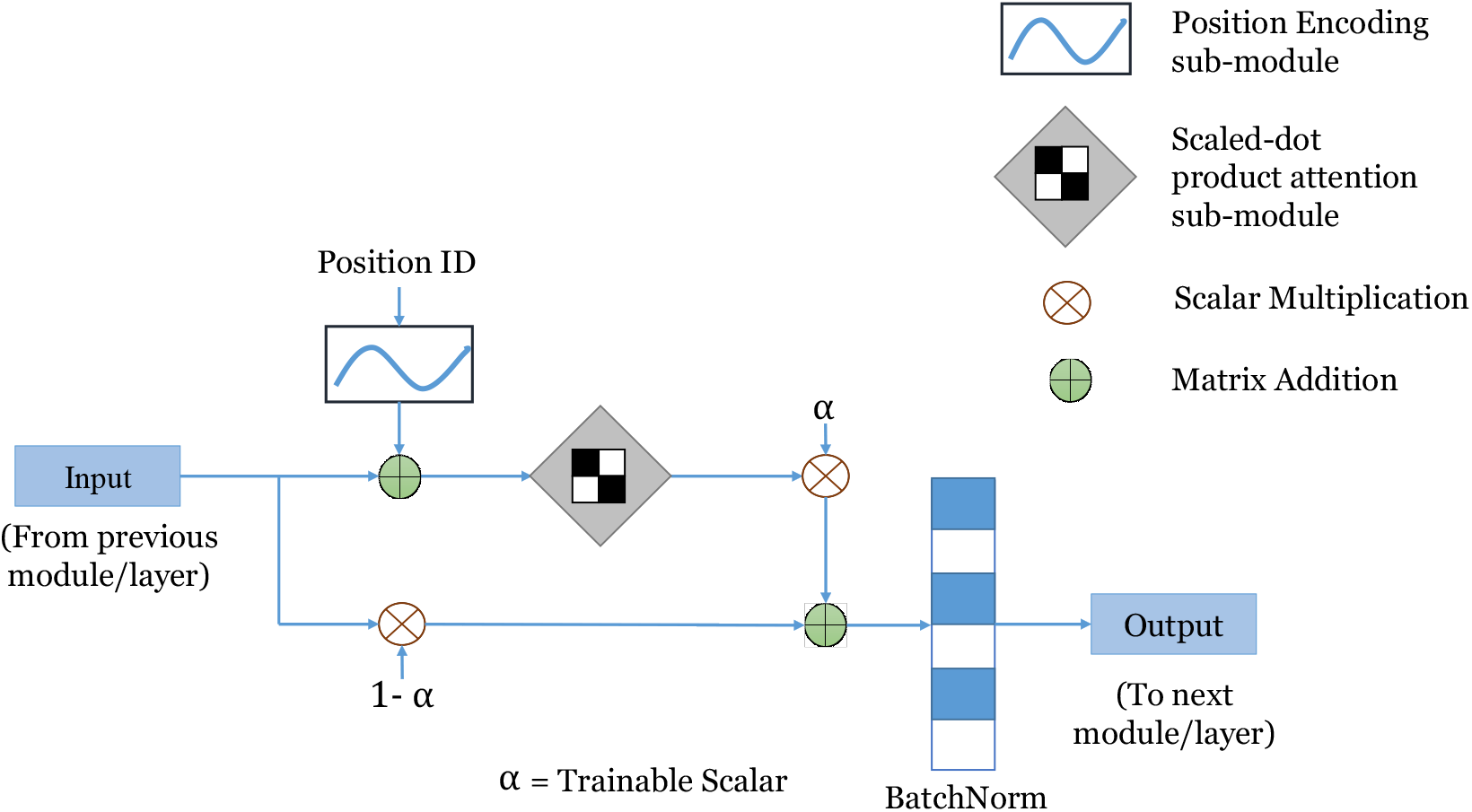
Architecture of the self-attention module used in SAINT.

##### Positional Encoding Sub-module

The objective of positional encodings is to inject some information about the relative or absolute positions of the residues in a protein sequence. The *Positional Encoding PosEnc_p_* for a position *p* can be defined as follows [55].

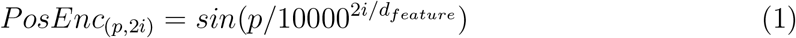

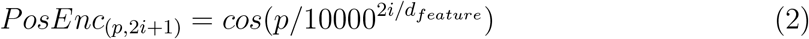

where *i* is the dimension. We used such function as it may allow the model to easily learn to attend by relative positions since for any fixed offset *k, PosEnc_p+k_* can be represented as a linear function of *PosEnc_p_* [55]. For every position *p, PosEnc_p_* has the dimension *d_pro_t_ein_* × *d_feature_*. The output of positional encoding is added with the inputs *x*, resulting in new representations *h* (see Eqn. 3) which contain not only the information extracted by the former layers or modules, but also the information about individual positions.

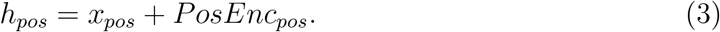

##### Scaled dot-product attention sub-module

The input features in this submodule, 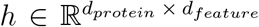 are first transformed into three feature spaces *Q, K* and *V*, representing *query, key* and *value* respectively, in order to compute the scaled dotproduct attention, where *Q*(*h*) = *Wçh, K* (*h*) = *W_K_ h, V* (*h*) = *W_V_ h*. Here *W_Q_, W_k_, W_V_* are parameter matrices to be learned. Figure 2 shows a schematic diagram of this module.

**Figure 2:**
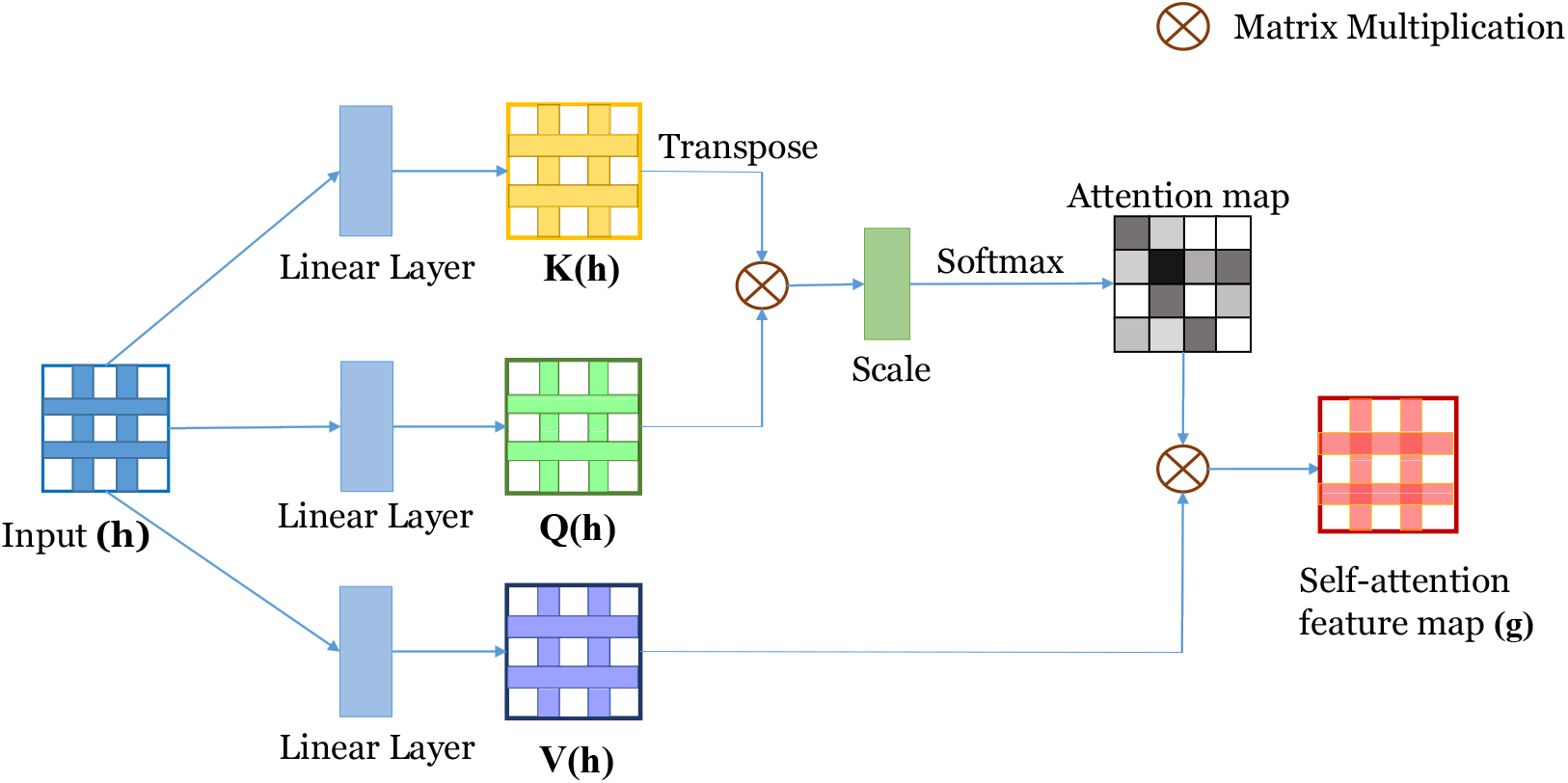
Architecture of the scaled dot-product attention sub-module.

Among various compatibility functions (e.g. scaled dot-product attention [55], additive attention [61], similarity-attention [64], multiplicative-attention [65], biased general attention [66], etc.), we have chosen the scaled dot-product attention as it showed much promise in case of sequential data. Vaswani *et al*. [55] showed that in practice, the dotproduct attention is much faster and space-efficient as it can be implemented using highly optimized matrix multiplication code, though theoretically both dot-product and additive attention have similar complexity. Scaled dot-product *S_i,j_* of two vectors *h_i_* and *h_j_* is calculated as shown in Equation 4.

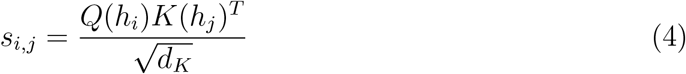

where *d_K_* is the dimension of the feature space *K*. The numerator of the equation, *Q*(*h_i_*)*K*(*h_j_*)*^T^* is the dot product between these two vectors, resulting in the similarity between them in a specific vector space. Here 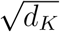 is the scaling factor which ensures that the result of the dot product does not get prohibitively large for very long sequences.

The attention weights *e* ∈ 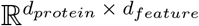 are calculated as shown in Equation 5, where *e_j,i_* represents how much attention have been given to the vector at position *i* while synthesizing the vector at position *j*.

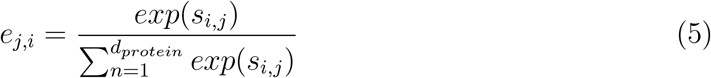

The attention distribution *e* is multiplied with the feature vectors *V*(*h*) and then in order to reduce the internal covariate shift this multiplicand is normalized using *batch normalization* [67], producing *g*, the output of the scaled dot-product attention submodule, following the Equation 6.

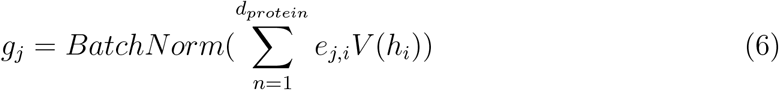

Here, *BatchNorm(.)* is the batch-normalization function and *g_j_* is the *j*-th vector in the output sequence of this sub-module. Finally, according to the Equation 7, *g* is multiplied by a scalar parameter *α,* the original input feature map *x* is multiplied by (1 – α) and these two multiplicands are summed to synthesize the final output *y*.

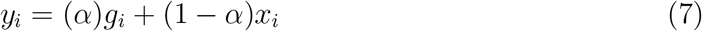

where *y_i_* is the *i*th output and *α* is a learnable scalar. By introducing weighed sum of *g_i_* and *x_i_*, we give our model the freedom to chose how much weight should be given to each of the features maps, *g_i_* and *x_i_* while generating the output *y_i_*. The optimal value of the parameter *α* is learnt through back propagation along with the rest of the model.

#### 2.2.2 Attention augmented inception-inside-inception (2A3I) module

A novel deep convolutional neural network architecture, *Inception*, was first introduced by Szegedy *et al*. [68], which demonstrated state-of-the-art performance for image classification and detection. An inception module has several branches, each having one or more convolutional layers. Fang *et al*. used an assembly of inception modules, which they call Inception-inside-Inception (3I module), in their proposed method MUFOLD-SS to predict protein secondary structure. They tried to leverage the inception blocks to retrieve both short-range and long-range dependencies and achieved the best known accuracy at the time. However, convolutional layers cannot capture enough information about long-range similarities or dependencies among feature vectors of a sequence, synthesized by a certain level of the network [69]. In protein secondary structure prediction, this issue leaves more impact on the overall accuracy when the sequence grows in length. Though these types of neural networks that use only convolutional layers need to be deeper to capture the long range dependency, it is often not feasible to add arbitrarily large numbers of layers. Moreover, the authors of MUFOLD-SS showed that using more than two inception-inside-inception modules sequentially does not result into significant increase in the overall accuracy, rather increases the computational expense. Earlier works [19,20,22,27,70,71] used Recurrent Neural Network(RNN) based architectures for capturing global features, but incorporating RNN or its derivatives (Gated Recurrent Units (GRU) [72], Long Short Term Memory (LSTM) [73]) inside 3I module would escalate the complexity and computational cost of the model. Therefore, we incorporated the self-attention mechanism to effectively capture both the short-range and long-range dependencies and to bring a better balance between the ability to model long-range dependencies and the computational efficiency. We placed our self attention modules in each branch of the 3I module as shown in Fig. 3. We call this an attention augmented inception-inside-inception (2A3I) module.

**Figure 3:**
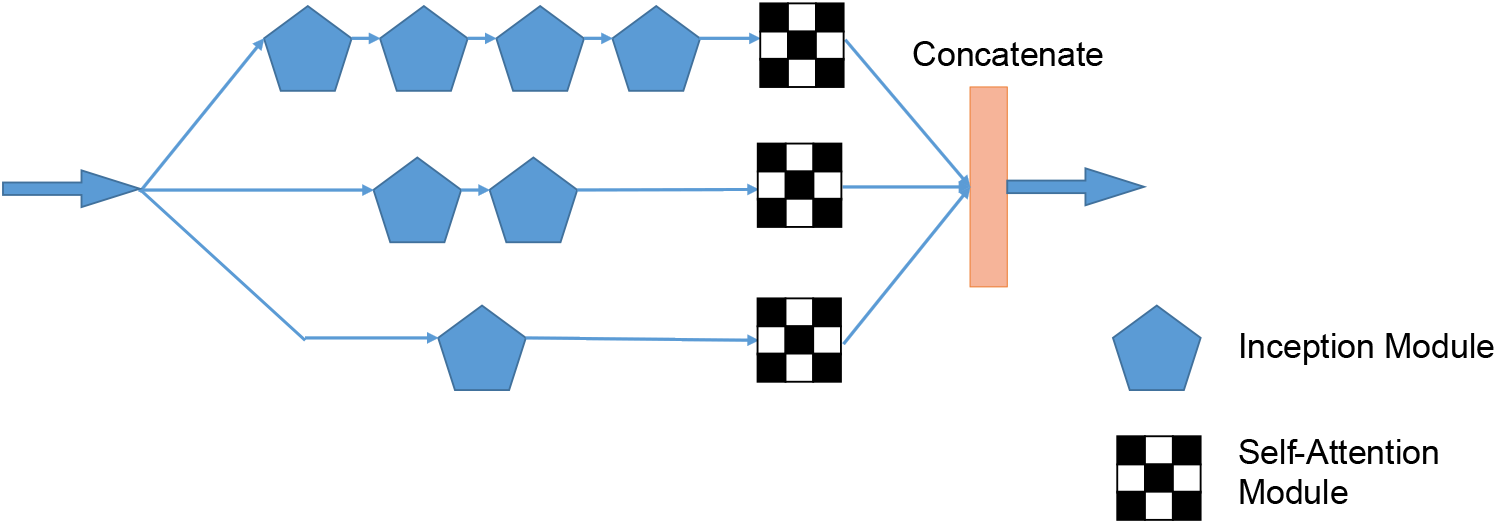
Architecture of our proposed 2A3I module by augmenting self-attention within the inception-inside-inception (3I) network.

#### 2.2.3 Overview of SAINT

A schematic diagram of the overall architecture of SAINT is depicted in Fig. 4. SAINT starts with two consecutive 2A3I modules followed by a self-attention module to supplement the non-local interactions captured by the initial two 2A3I modules. We also observed that this attention module helps achieve faster learning rate. MUFOLD-SS used one convolutional layer with window size 11 after two 3I modules. The level of long-range interactions being captured varies with varying lengths of the window. However, we observed that using window size larger than 11 increases the computational cost without significantly increasing the performance. As a result, we used similar convolutional layer as MUFOLD-SS. However, we included another self-attention module after the convolutional layer to help capture the relations among vectors that the convolutional layer failed to retrieve. The last two dense layers in the MUFOLD-SS were also used in SAINT. However, we placed an attention module in between the two dense layers. We did so to understand how the residues align and interact with each other just before generating the output. This paves the way to have an interpretable deep learning model (as we will discuss in Sec. 3.3.1).

**Figure 4:**
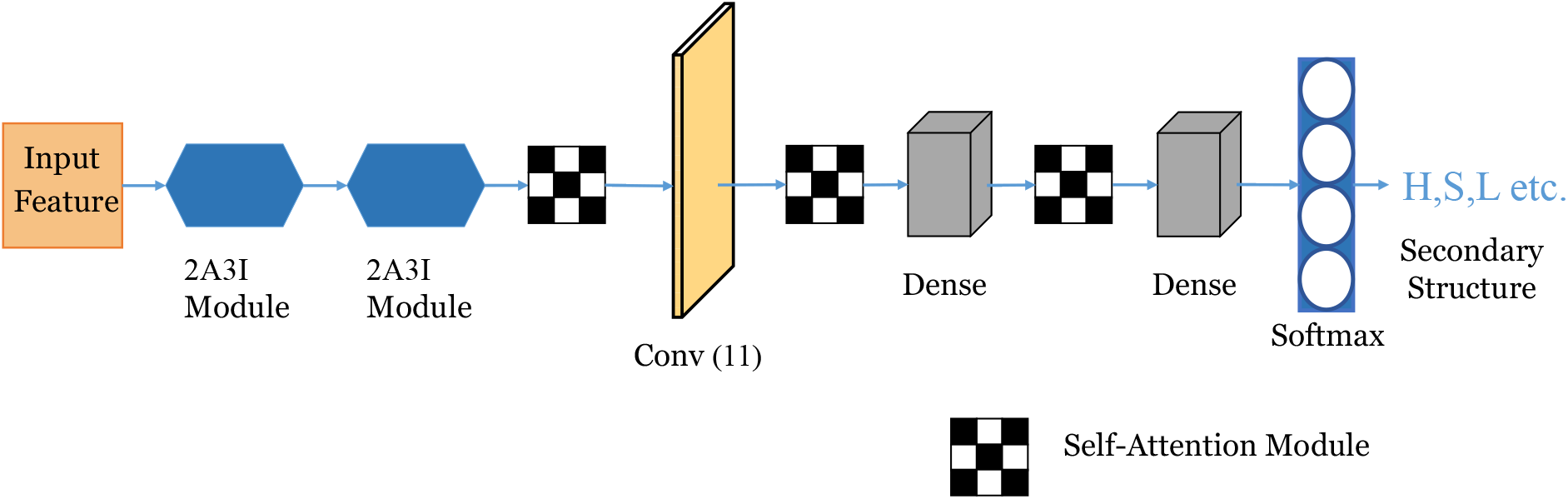
A schematic diagram of the overall architecture of SAINT. It comprises two 2A3I modules, three self-attention modules, convolutional layers with window size 11 and two dense layers.

## 3 Results and Discussion

We performed an extensive evaluation study, comparing SAINT with the state-of-the-art Q8 prediction methods on a collection of publicly available benchmark datasets.

### 3.1 Dataset

To make a fair comparison with the most recent state-of-the-art methods, especially with SPOT-1D (the most accurate method to date), we used the same training and validation sets that were used in SPOT-1D. However, apart from comparing our model against the most recent and large test sets *TEST2016* and *TEST2018* generated and analyzed by Hanson *et al*. [44,60], we evaluated SAINT on CASP12, CASP13 and the template free modelling targets of four CASP datasets (CASP10 ~ CASP13). CB513 [41], which is a relatively old, yet widely used benchmark dataset has been excluded from our evaluation as there are many sequences in CB513 with > 25% sequence similarity to the training set.

The training set contains 10,029 proteins from CullPDB with resolution < 2.5 Å, R-factor < 1.0 and a sequence identity cutoff of 25% according to BlastClust (Altschul *et al*. [56]). The validation set contains 983 proteins from CullPDB with the same specifications applied to training set (see [44] and [60] for more details on these dataset). We provide brief descriptions of the test sets in subsequent sections.

#### 3.1.1 TEST2016

TEST2016 dataset contains 1,213 proteins that were all deposited on PDB between June 2015 and February 2017 with similar parameter settings as the training set and do not contain more than 700 residues. It has less than 25% sequence similarity with the training and validation sets according to BlastClust [56]. It was compiled by Hanson *et al*. [60] and is available at https://servers.sparks-lab.org/downloads/SPOT-1D-dataset.tar.gz.

#### 3.1.2 TEST2018

TEST2018 dataset contains 250 high-quality, non-redundant proteins that were all deposited on PDB between January 2018 and July 2018. The dataset was also filtered to remove redundancy at a 25% sequence identity cutoff and to remove proteins having more than 700 residues. It was generated by Hanson *et al*. [60] and is available at https://servers.sparks-lab.org/downloads/SPOT-1D-dataset.tar.gz.

#### 3.1.3 CASP

CASP stands for Critical Assessment of protein Structure Prediction. This is an biennial competition for protein structure prediction and a community wide effort to advance the state-of-the-art in modelling protein structure from its amino acid sequences since 1994 [74]. Among the CASP datasets, we took into account the most recent ones CASP13 and CASP12. We removed one domain sequence out of 32 in CASP13 (T0951-D1) and six domain sequences out of 55 in CASP12 as they had more than 25% sequence similarity to the training set according to CD-HIT [75]. Apart from these, we prepared a dataset comprising the template free modelling (FM) targets in CASP datasets to show the performance of SAINT where the query sequences do not have statistically significant similar protein sequences with known structures. Some of the FM targets had *>* 25% sequence similarity with our training set, which were therefore excluded from the test set. Thus, we compiled a test set which we call *CASP-FM* comprising 56 domain sequences: 10 FM targets from CASP13, 22 FM targets from CASP12, 16 FM targets (out of 30) from CASP11, and 8 FM targets (out of 12) from CASP10. The CASP proteins were downloaded from its official website http://predictioncenter.org/.

### 3.2 Method comparison

We compared SAINT with the most recent and accurate Q8 predictors: MUFOLD-SS [30], NetSurfP [43] and SPOT-1D [44]. These state-of-the-art methods have been shown to outperform other popular Q8 predictors, namely, SSPro8 [19], RaptorX-SS8 [40], DeepGSN [26], DeepCNF [25], DCRNN [27], NCCNN [28], CNNH_PSS [29], CBRNN [71], etc.

We evaluated the methods under various evaluation metrics, such as, Q8 accuracy, precision, recall and *F*1-score. We performed Wilcoxon signed-rank test (with *α* = 0.05) to measure the statistical significance of the differences between SAINT and each of the compared state-of-the-art methods.

### 3.3 Results on benchmark dataset

The comparison of SAINT with the state-of-the-art Q8 structure prediction methods on TEST2016, TEST2018, CASP13, CASP12, and CASPFM is shown in Table 1. To train SAINT and tune necessary hyper-parameters, we have used the same training and validation sets that were used by SPOT-1D. Notably, SPOT-1D is an ensemble of 9 models where each single model uses predicted contact map in addition to other features. SAINT, on the other hand, is an ensemble of only 4 models, 3 of which take advantage of predicted contact map with different windows sizes. Experimental results show that SAINT outperforms all other methods across all the test sets. It is worth mentioning that SAINT’s accuracy on the validation set (78.18%) was also better than that of SPOT-1D (77.60%). SPOT-1D’s base model, which does not require contact maps as features is also an ensemble of 9 models, whereas SAINT-base model is a single model. Despite being a single model, SAINT-base consistently outperformed SPOT-1D base in TEST2016 and TEST2018. We could not evaluate SPOT-1D base on CASP12, CASP13, and CASPFM as it is not publicly available. From Table 1, it is also evident that SAINT is substantially better than the other recent methods, namely, NetSurfP-2.0 and MUFOLD-SS. Even the base model of SAINT consistently outperformed both NetSurfP-2.0 and MUFOLD-SS. The remarkably large improvement of SAINT over MUFOLD-SS across all the dataset suggests the advantage of augmenting our proposed self-attention mechanism in the Deep3I network used in MUFOLD-SS. Statistical tests (see Table 2) suggests that these improvements of SAINT over other methods are statistically significant (*P* << 0.05).

**Table 1:** A comparison of the Q8 accuracy (%) obtained by SAINT and other state-of-the-art methods on TEST2016 and TEST2018 dataset. Best results for each benchmark dataset are shown in bold.

| Method | TEST2016 | TEST2018 | CASP13 | CASP12 | CASPFM |
| --- | --- | --- | --- | --- | --- |
| SAINT<sup>1</sup> | **77.73** | **76.10** | **74.78** | **74.17** | **72.25** |
| SPOT-1D<sup>1</sup> | 77.10 | 75.41 | 73.91 | 73.67 | 71.85 |
| SAINT-base | 76.23 | 74.48 | 73.5 | 71.78 | 70.00 |
| SPOT-1D-base<sup>1,2</sup> | 76.03 | 74.26 | - | - | - |
| NetSurfP-2.0 | 75.68 | 73.04 | 72.85 | 71.43 | 70.16 |
| MUFOLD-SS | 75.56 | 73.66 | 73.44 | 71.44 | 70.21 |

<sup>1</sup> Indicates ensemble model
<sup>2</sup> Not publicly available. Results reported by SPOT-1D [44].

**Table 2:** Statistical significance of the Q8 accuracy between SAINT and other state-of-the-art methods. The numbers of protein chains or domains in these datasets are shown in parentheses. We show the *p*-values using a Wilcoxon signed rank test.

| Method | TEST2016<br>(1213) | TEST2018<br>(250) | CASP13<br>(31) | CASP12<br>(49) | CASPFM<br>(56) |
| --- | --- | --- | --- | --- | --- |
| SPOT-1D | 8.168e-27 | 3.893e-5 | 0.101 | 0.0345 | 0.0791 |
| NetSurfP-2.0 | 2.607e-57 | 3.258e-18 | 0.179 | 1.55e-6 | 0.0001 |
| MUFOLD-SS | 1.531e-88 | 3.145e-21 | 0.179 | 6.51e-5 | 0.005 |

In addition to the model accuracy, we also investigated the *precision, recall* and *F*1-score to obtain better insights on the performances of various methods. Precision, also know as predictivity, denotes the confidence that can be imposed on a prediction. Recall signifies how accurately an algorithm can predict a sample from a particular class. Sometimes an algorithm tends to over-classify which results into high recall but low precision. On the other hand, some algorithms tend to under-classify, preserving the precision at the cost of recall. In order to get an unbiased evaluation of the performance, *F*1-score is considered to be an appropriate measure and has been being used for over 25 years in various domains [76,77]. Tables 3, 4, and 5 show the precision, recall and *F*1-score on each of the 8 states obtained by SAINT and other methods. These results suggest that SAINT achieves better *F*1-score than other methods on 5 states (out of 8 states), showing that SAINT produced more balanced and meaningful results than other methods. SAINT substantially outperforms other methods on the non-ordinary states [25] such as I, G, S, and T. However, MUFOLD-SS and SPOT-1D achieved slightly better *F*1-score for the ‘B’ and ‘E’ states respectively. State ‘I’ (*π*-helix) is extremely rare which comprises seven or more residues and is present in 15% of all known protein structures [78]). They are very difficult to predict, but mostly found at functionally important regions such as ligand- and ion-binding sites [78]. Therefore, specialized predictors, such as PiPred [78], is also available that only predicts the *π*-helix structures. SAINT significantly outperforms SPOT-1D, NetSurfP-2.0, and MUFOLD-SS in predicting *π*-helix in TEST2016 dataset by correctly predicting 21 out of 47 ‘I’ states and thus achieving a recall of 0.45 for this structure. SAINT’s precision for *π*-helix, on the other hand, is 1. This is remarkable considering the fact that the *π*-helix specific predictor, PiPred, reports precision and recall of 0.48 and 0.46 respectively on a different dataset which they analyzed in [78].

**Table 3:** Predictive precision on each of the 8 states obtained by SAINT and other state-of-the-art methods on TEST2016 dataset.

| Q8 Label | SAINT | SPOT-1D | NetSurfP-2.0 | MUFOLD-SS |
| --- | --- | --- | --- | --- |
| H | 0.879 | 0.884 | **0.885** | 0.868 |
| B | **0.76** | 0.671 | 0.65 | 0.609 |
| E | 0.843 | **0.852** | 0.822 | 0.85 |
| G | **0.581** | 0.547 | 0.536 | 0.519 |
| I | **1** | 1 | 0.044 | 0.857 |
| T | **0.663** | 0.641 | 0.615 | 0.631 |
| S | **0.639** | 0.624 | 0.579 | 0.589 |
| C | **0.648** | 0.631 | 0.613 | 0.607 |

**Table 4:**
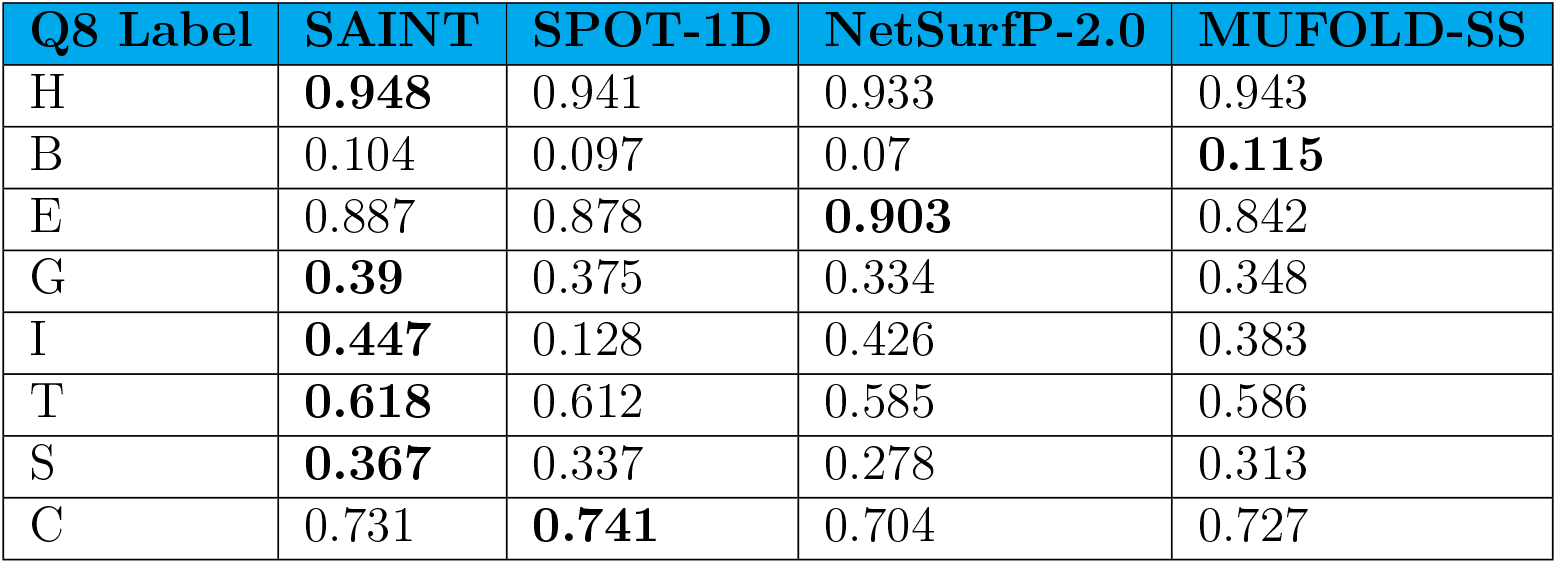
Recall on each of the 8 states obtained by SAINT and other state-of-the-art methods on TEST2016 dataset.

**Table 5:** *F*1-score on each of the 8 states obtained by SAINT and other state-of-the-art methods on TEST2016 dataset.

| Q8 Label | Frequency | SAINT | SPOT-1D | NetSurfP-2.0 | MUFOLD-SS |
| --- | --- | --- | --- | --- | --- |
| H | 98139 | **0.912** | 0.911 | 0.908 | 0.904 |
| B | 3018 | 0.183 | 0.169 | 0.126 | **0.193** |
| E | 62657 | 0.864 | **0.865** | 0.861 | 0.846 |
| G | 10770 | **0.467** | 0.445 | 0.412 | 0.417 |
| I | 47 | **0.618** | 0.227 | 0.079 | 0.529 |
| T | 32297 | **0.639** | 0.626 | 0.599 | 0.608 |
| S | 23466 | **0.466** | 0.438 | 0.376 | 0.409 |
| C | 57483 | **0.687** | 0.682 | 0.655 | 0.662 |

We analyzed the CASPFM dataset comprising the free modeling targets in the CASP dataset to demonstrate the performance of models on proteins with previously unseen folds. SAINT achieved the best accuracy on CASPFM, suggesting SAINT’s superiority in predicting structures of proteins having very low sequence homology with proteins of known structures.

While the advantage of utilizing our proposed self-attention mechanism in the Deep3I framework of MUFOLD-SS is evident from the significant improvement of SAINT over MUFOLD-SS across all the dataset analyzed in this study, we further investigated the efficacy of our proposed attention mechanism in capturing the long-range interactions. We computed the number of non-local interactions per residue for each of the 1,213 proteins in TEST2016, and sorted them in an ascending order. Next, we put them in six equal sized bins *b_1_,b_2_,…, b_6_* (each containing 202 proteins except for *b_6_* which contains 203 proteins), where b_1_ contains the proteins with the lowest level of non-local interactions and *b_6_* represents the model condition with the highest level of non-local interactions. We show the Q8 accuracy of SAINT-base and MUFOLD-SS on these model conditions in Table 6 and Fig. 5. Note that, instead of our ensemble model which is more accurate than our base model, we deliberately show the results for our single base model, which uses the same feature set as MUFOLD-SS, and the only difference between them is the self-attention modules introduced in our architecture. These results show that the difference in predictive performance between SAINT-base and MUFOLD-SS significantly increases with increasing levels of non-local interactions. There is no statistically significant difference between them on *b***i**, but as we increase the level of non-local interactions, SAINT becomes significantly more accurate than MUFOLD-SS and attains the highest level of improvement on *b_6_*. This clearly indicates that capturing non-local interactions by self-attention is the key factor in the improvement. We also performed the same analyses on other methods (see Fig. 6). The results in Fig. 6 show that the differences among of these methods are not that substantial on the model conditions with low levels of long-range interactions, but the differences become notable as we increase the non-local interactions. SAINT not only achieved the best accuracy, its improvement over other methods increases with increasing amount of long-range interactions as well – suggesting the superiority of our proposed self-attention mechanism compared to CNN+LSTM (used in NetSurfP-2.0) and CNN (ResNet)+BRLSTM (used in SPOT-1D) in terms of capturing the non-local interactions.

**Figure 5:**
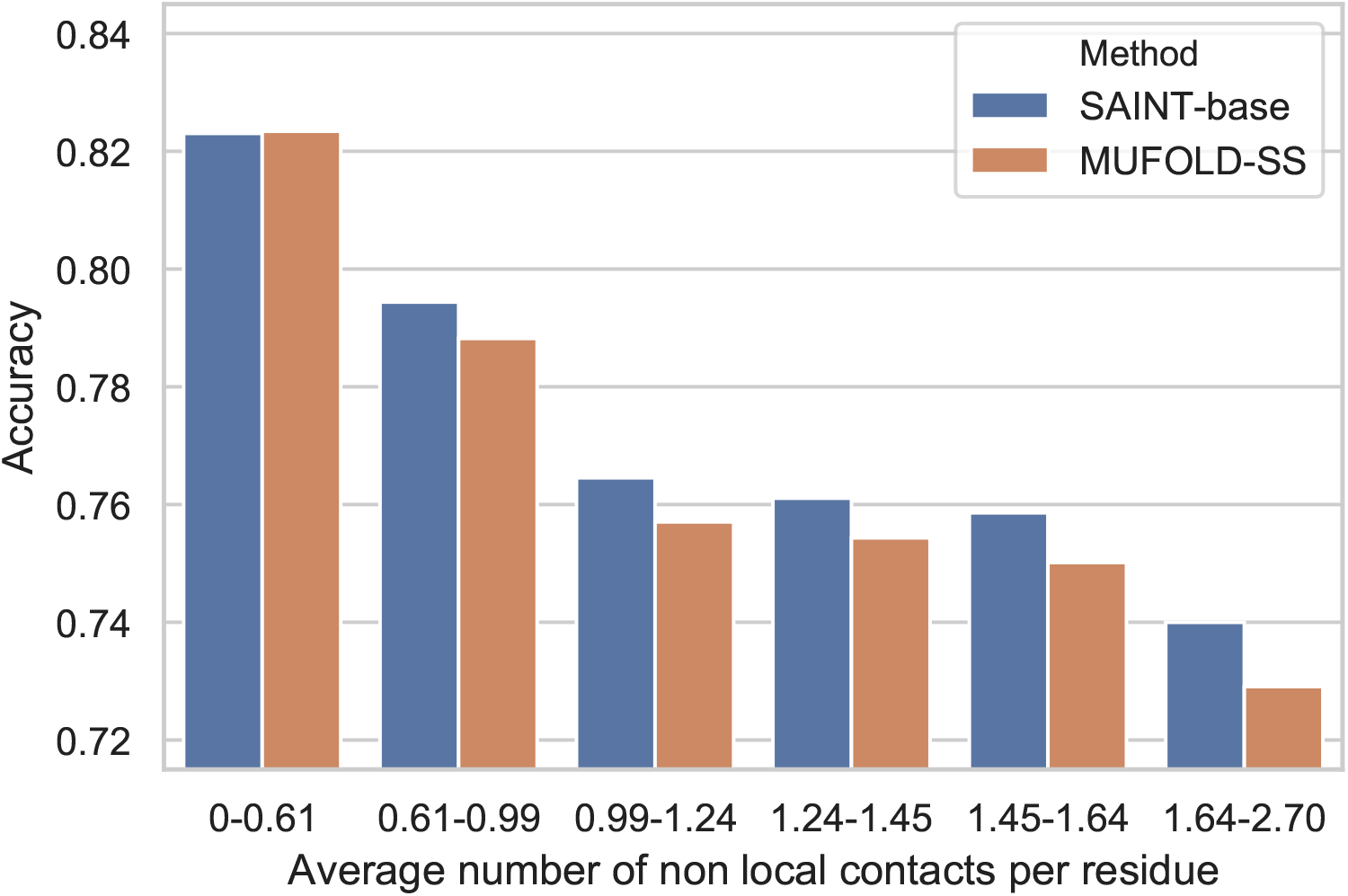
Accuracy of SAINT-base and MUFOLD-SS under various levels of non-local interactions. We show the results on the TEST2016 test set using six bins of proteins as shown in Table 6.

**Figure 6:**
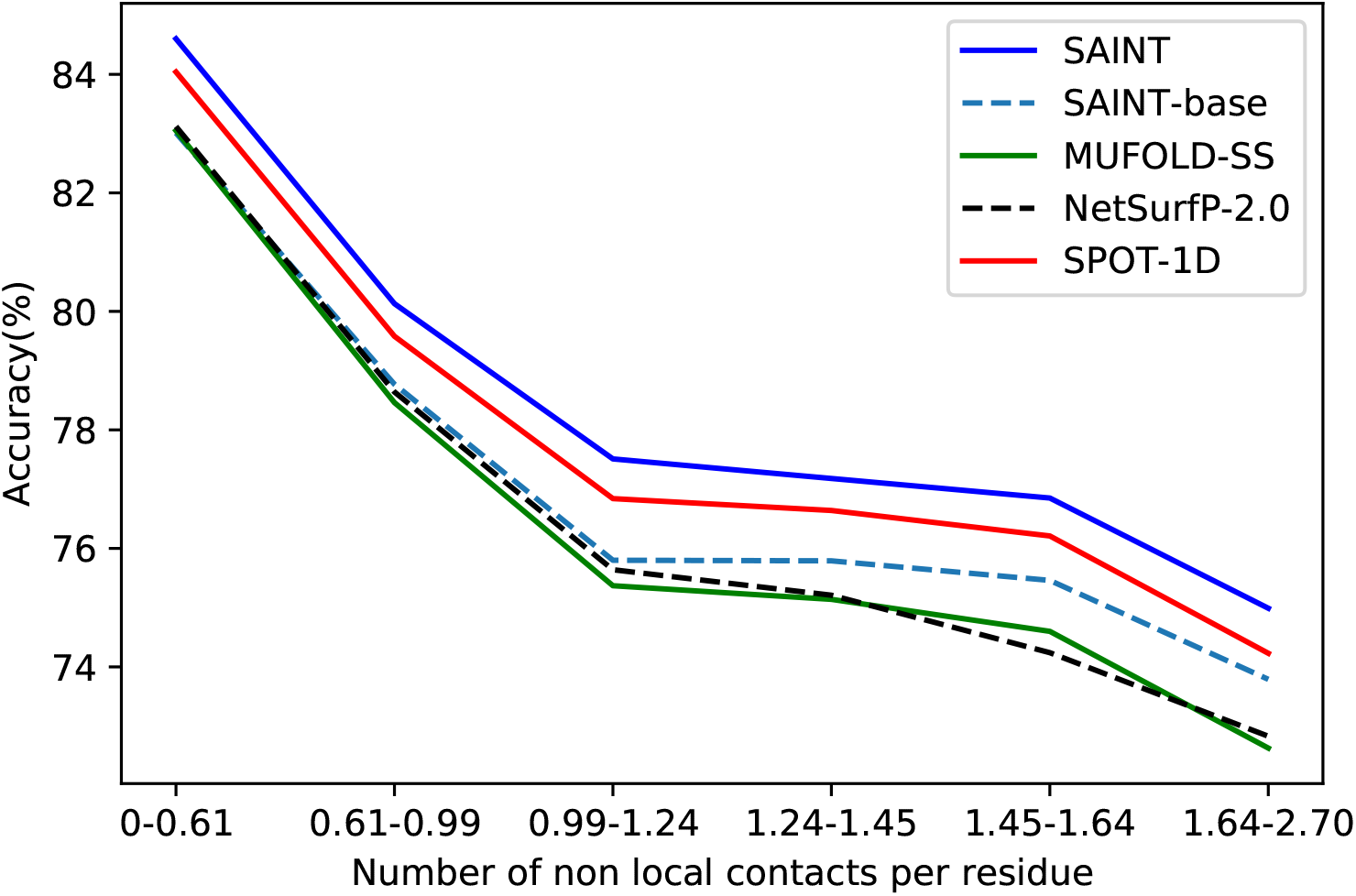
Accuracy of SAINT, SPOT-1D, NetSurfP-2.0 and MUFOLD-SS as a function of the average number of non-local interactions per residue. We show the results on the six bins as shown in Table 6.

**Table 6:** Accuracy of SAINT(base) and MUFOLD-SS under various levels of non-local interactions. 1,213 proteins in TEST2016 were divided into 6 disjoint bins each having 202 proteins except the last one which had 203 proteins. The binning was based on the number of non-local contacts per residue in the proteins.

| Non-local contacts per residue | Q8 Accuracy (%) of SAINT-base | Q8 Accuracy (%) of MUFOLD-SS | Accuracy Difference | $p$-value |
| --- | --- | --- | --- | --- |
| 0-0.61 | 83.01 | 83.05 | -0.04 | 0.721287 |
| 0.61-0.99 | 78.77 | 78.46 | 0.31 | 0.01036 |
| 0.99-1.24 | 75.80 | 75.37 | 0.43 | 0.004 |
| 1.24-1.45 | 75.79 | 75.14 | 0.65 | 0.0003 |
| 1.45-1.64 | 75.46 | 74.60 | 0.86 | 0.0001 |
| 1.64-2.70 | 73.79 | 72.63 | 1.16 | 1.36e-6 |

In order to demonstrate the efficacy of SAINT and other methods in capturing the continuous structure of a protein, we show the one-dimensional map of the native and predicted secondary structure of a representative protein 5M2PA in TEST2016 (see Fig. 7).

**Figure 7:**
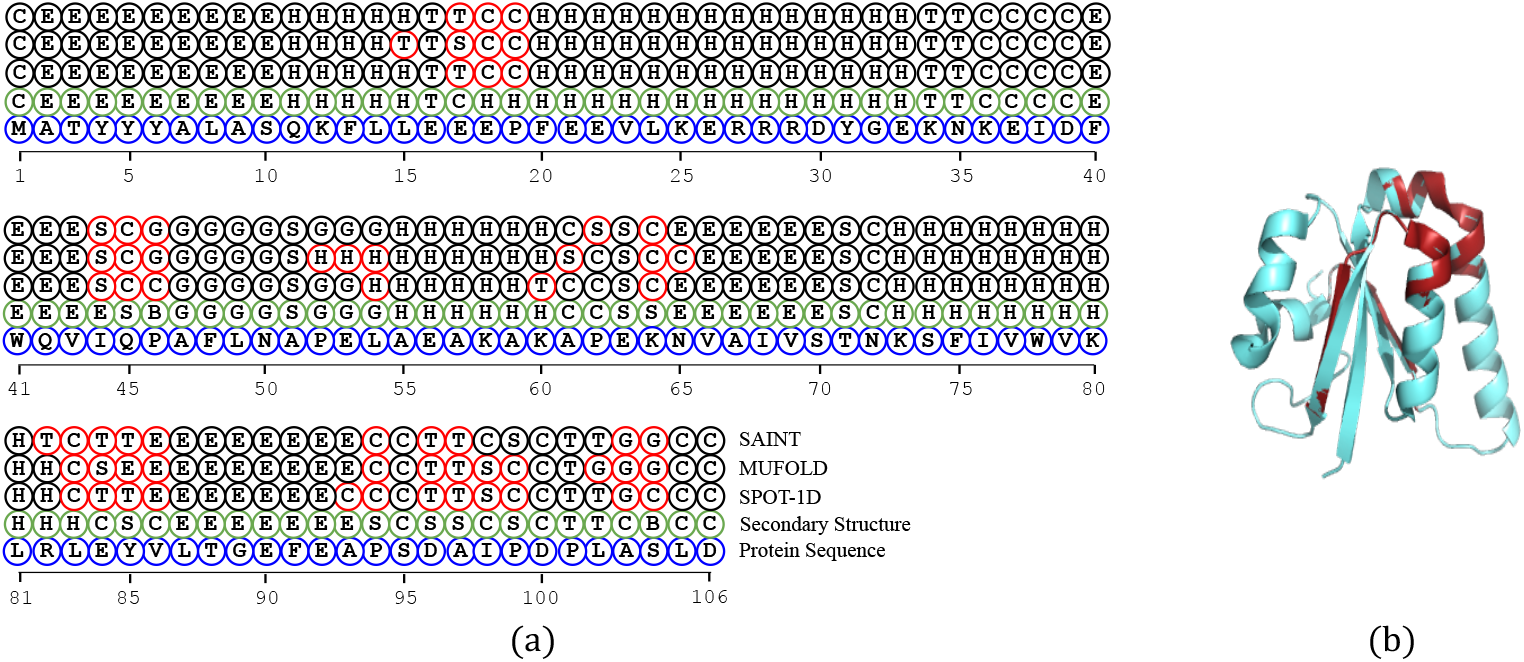
Structure prediction on 5M2PA protein chain by various methods. (a) One-dimensional map of the native structure of 5M2PA and the predicted structures by various methods. (b) Superposition of the structures predicted by SAINT (cyan) with the structures obtained from PDB (red) for 5M2PA. This image is generated in Pymol [98]. Since Pymol does not differentiate between all the 8 distinct states, we translated the 8-state structure to 3-state structure according to the Rost and Sander scheme [10].

#### 3.3.1 Interpretability

One notable feature of SAINT is that, unlike most of the existing deep learning techniques, it can provide insights on how the architecture is making decisions, especially regarding the long-range interactions. Self-attention alignment matrix has already been used to interpret how different parts of input are dependent on each other while generating the output [55, 79–82], and hence it was used to develop interpretable models [83–86]. Thus, we made an attempt at using attention matrix to capture and provide insight into the long-range interactions. Long-range interactions are crucial for predicting the secondary structure of proteins. For example, a secondary structure state *β*-strand is stabilized by hydrogen bonds formed with other *β*-strands that can be far apart from each other in the protein sequence [26].

As mentioned earlier, we placed an attention module just before the last dense layer in the architecture of SAINT. In addition to improving the prediction performance, a motivation behind this attention module has been to introduce some form of interpretability to the deep learning model. Indeed we are able to relate the self-attention alignment score matrix of this attention module to the spatial proximity of a residue with other residues far apart in the primary sequence. In Fig. 8, we show the relation between the spatial proximity and attention scores for a sample protein 5epmD in TEST2016. We selected a short sequence 5epmD (only 33 residues) to easily demonstrate with visualizations how the alignment matrix provides insight about the long-range interactions. We show the distances of the first five residues (‘D’, ‘C’, ‘L’, ‘G’, ‘M’) to all other subsequent residues as line graphs and superimpose them on the attention matrix obtained by a single model of SAINT. We choose only the first 5 residues for the sake of readability and clarity of this figure. For this protein, highest attention has been given to the 15-th residue ‘K’ and 28-th residue ‘W’, meaning that most of the residues generated the highest attention score with respect to these two residues. Interestingly, these two residues are where the spatial distance lines reach their local minima, indicating a possible turn, bend or contact pair. Being inspired by this, we systematically analyzed the attention matrices and spatial distance graphs of all the 1213 proteins in TEST2016. We consider only those “downslopes” in the spatial proximity line graphs that continue to decrease for at least three consecutive residues, and thereby ignoring very small decreasing regions which span less than 3 residues. We have observed that, in 93.33% of these proteins, the residues with most attention on them are within a downslope region of another residue’s spatial distance line curve. This indicates that the residues with relatively higher attention scores are likely to be spatially closer to some other distant residues in the primary sequence. While these results are promising, especially considering the black-box nature of other deep learning based methods, they should be interpreted with care. The long-range interactions suggested by the attention matrix may contain false positives and false negatives. Higher attention scores do not necessarily guarantee a contact pair, nor is it certain that all the contact pairs will have relatively higher attention scores. More work is required to design an attention mechanism so that the attention matrix is more closely related to the contact map. This is an interesting research avenue which we left as a future work. We believe that this matrix with appropriate modifications will be useful to understand the complex relationship between the primary sequence and various structural and functional properties of proteins.

**Figure 8:**
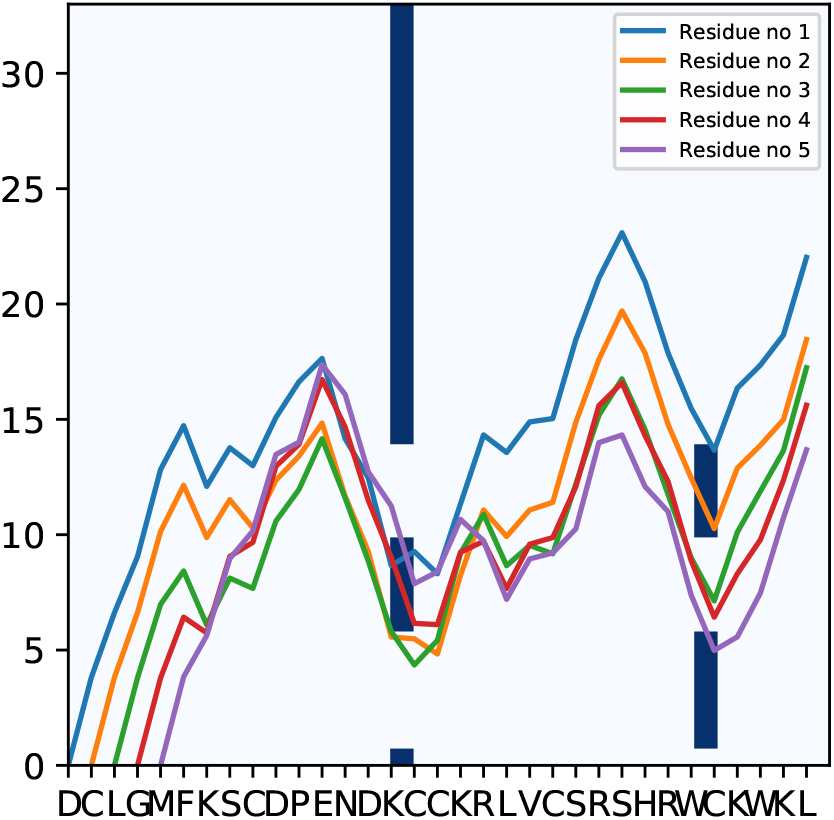
Demonstration of the interpretability of SAINT using the attention map. Spatial distances of the first five residues (‘D’, ‘C’, ‘L’, ‘G’, ‘M’) in 5epmD to all other subsequent residues are shown by line graphs and they are superimposed on the attention matrix. Deeper hue on the 15-th residue ‘K’ and the 28-th residue ‘W’ indicates higher attention scores.

#### 3.3.2 Running time

SAINT is much faster than the best alternate method SPOT-1D. For generating the structures of 1,213 protein chains in TEST2016, given the necessary input files, SAINT took approximately 360 ± 5 seconds whereas SPOT-1D took approximately 2, 485 ± 5 seconds on our local machine (Intel corei7-7700 CPU 3.60 GHz (4 cores), 16GB RAM, NVIDIA GeForce GTX 1070 GPU). Under the same settings, SAINT took approximately 197 ± 5 seconds to generate secondary structures for the 250 proteins in TEST2018, whereas SPOT-1D took approximately 668 ± 5 seconds. Since both these methods use the same input files for feature generation, this substantial difference in running time can be attributed to the efficiency of our attention based method over the LSTM networkbased model used in SPOT-1D.

## 4 Conclusions

We have presented SAINT, a highly accurate, fast, and interpretable method for 8-state SS prediction. We demonstrate for the first time that the self-attention mechanism proposed by Vaswani *et al*. [55] is a valuable tool to apply in the structural analyses of proteins. Another earlier type of attention mechanism proposed by Bahdanau *et al*. [61] coupled with recurrent neural network (RNN) based encoder-decoder architectures achieved state-of-the-art performance on various natural language processing tasks (e.g. neural machine translation [65,87], question answering task [88,89], text summarization [90,91], document classification [92,93], sentiment classification [94,95], etc.). As proteins are also sequences similar to sentences in a language, this type of architecture is expected do well in protein secondary structure prediction as well. However, previous attempts [96] on using attention with LSTM based encoder-decoder only achieved 68.4% accuracy on CB513 dataset which is significantly worse than the performance of MUFOLD-SS (70.63% on CB513) [30]. In this study, we have used the self-attention mechanism in a unique way and proposed a novel attention augmented 3I module (2A3I module) and achieved notable success. We have used the self-attention mechanism to retrieve the relation between vectors that lay far from each other in a sequence. As self-attention mechanism looks at a single vector and measures its similarity or relationship with all other vectors in the same sequence, it does not need to encode all the information in a sequence into a single vector like recurrent neural networks. This reduces the loss of contextual information for long sequences.

SAINT contributes towards simultaneously capturing the local and non-local dependencies among the amino acid residues. Unlike some of the existing deep learning methods, SAINT can capture the long-range dependencies without using computationally expensive recurrent networks or convolution networks with large window sizes. SAINT was assessed for its performance against the state-of-the-art 8-state SS prediction methods on a collection of widely used benchmark dataset. Experimental results suggest that SAINT consistently improved upon the best existing methods across various widely used benchmark dataset.

One of the most significant conclusions from the demonstrated experimental results is that appropriate use of self-attention mechanism can significantly boost the performance of deep neural networks and is capable of producing results which rank SAINT at the very top of the current SS prediction methods. Thus, the idea of applying self-attention mechanism can be applied to predicting various other protein attributes (e.g., torsion angles, turns, etc. [97]) as well. Therefore, we believe SAINT advances the state-of-the-art in this domain, and will be considered as a useful tool for predicting the secondary structures of proteins.

